# Seasonal and spatial variation in growth and abundance of zebra mussel (*Dreissena polymorpha*) in a recently invaded lake: implications for management

**DOI:** 10.1101/656371

**Authors:** Matteo Rolla, Sonia Consuegra, David J. Hall, Carlos Garcia de Leaniz

## Abstract

The control of the highly invasive zebra mussel (*Dreissena polymorpha*) has been flagged as a priority but success has been variable. A better understanding of the growth and drivers of settlement of zebra mussel is necessary for a more efficient management of this invasive species, but seasonal data are still relatively scant. We monitored the seasonal changes in settlement rates, density, and growth of zebra mussel in artificial substrates over one year in Cardiff Bay (UK), an artificial amenity lake invaded by zebra mussels in 2003 and where the species is rapidly expanding. Mean settling rates varied from 4,200 to 6,200 mussel m^−2^ over June to September mirroring changes in water temperature, and peaked at 17,960 mussels m^−2^, the highest density reported in Britain. Density was highest at the deepest panels (3 m). Growth varied significantly among sampling stations, with growth taking place during the summer and ceasing during winter and spring. Mixture analysis reveals the existence of multiple cohorts displaying different growth and settling rates and that follow different density dependent mechanisms, being positive density-dependent at low densities, and negative density-dependent at high densities. We suggest this creates the conditions necessary for source and sink metapopulations to develop which may need to be taken into account in management. Targeting mussels for removal in deep waters during the summer and early autumn might prove beneficial, but the existence of contrasting density-dependent mechanisms suggests that removal may be beneficial or counterproductive depending on local conditions.

## Introduction

The zebra mussel (*Dreissena polymorpha*) is one of the most damaging aquatic invaders (Strayer 2010), having been included in the list of the 100 world’s worst alien species (Lowe et al. 2000). Zebra mussels can drastically reduce the biomass of phytoplankton, and change its community composition, which can in turn change water parameters, resuspend nutrients into the water column (Bastviken et al. 1998), and increase water transparency (Fahnenstiel et al. 1995; Holland 1993). The species can compete for food and space with native freshwater mussels and drive them to extinction through epizootic colonization, disruption of their valve functionality, smothering of their siphons, and impairing of their movement through deposition of metabolic waste (Baker & Hornbach 1997; Schloesser et al. 1996). A strong association has been found between density of zebra mussels and mortality of native unionid mussels (Ricciardi et al. 1995).

The economic losses caused by zebra mussels can be considerable. They can block pipes and water supplies, intakes from nuclear, hydroelectric and industrial facilities (O’Neill 1997), and clog the cooling systems of power boats (Johnson et al. 2001). Navigation buoys have been sunk under the weight of zebra mussels, and dock pilings can become severely weakened due to zebra mussel encrustations (Minchin & Moriarty 1998). In the US, the cost of cleaning a single hydroelectric plant of zebra mussels may amount to $92,000 per year, and the combined costs may have reached $6.5 billion over 10 years (Lovell et al. 2006). Not surprisingly, the control of zebra mussel has become a priority (Aldridge et al. 2004; Aldridge et al. 2006; Strayer 2010). However, eradication measures are not always successful (Lund et al. 2018; Whitledge et al. 2015), may require several treatments, and may have to be extended over several years (Table 1.).

**Table 1.**
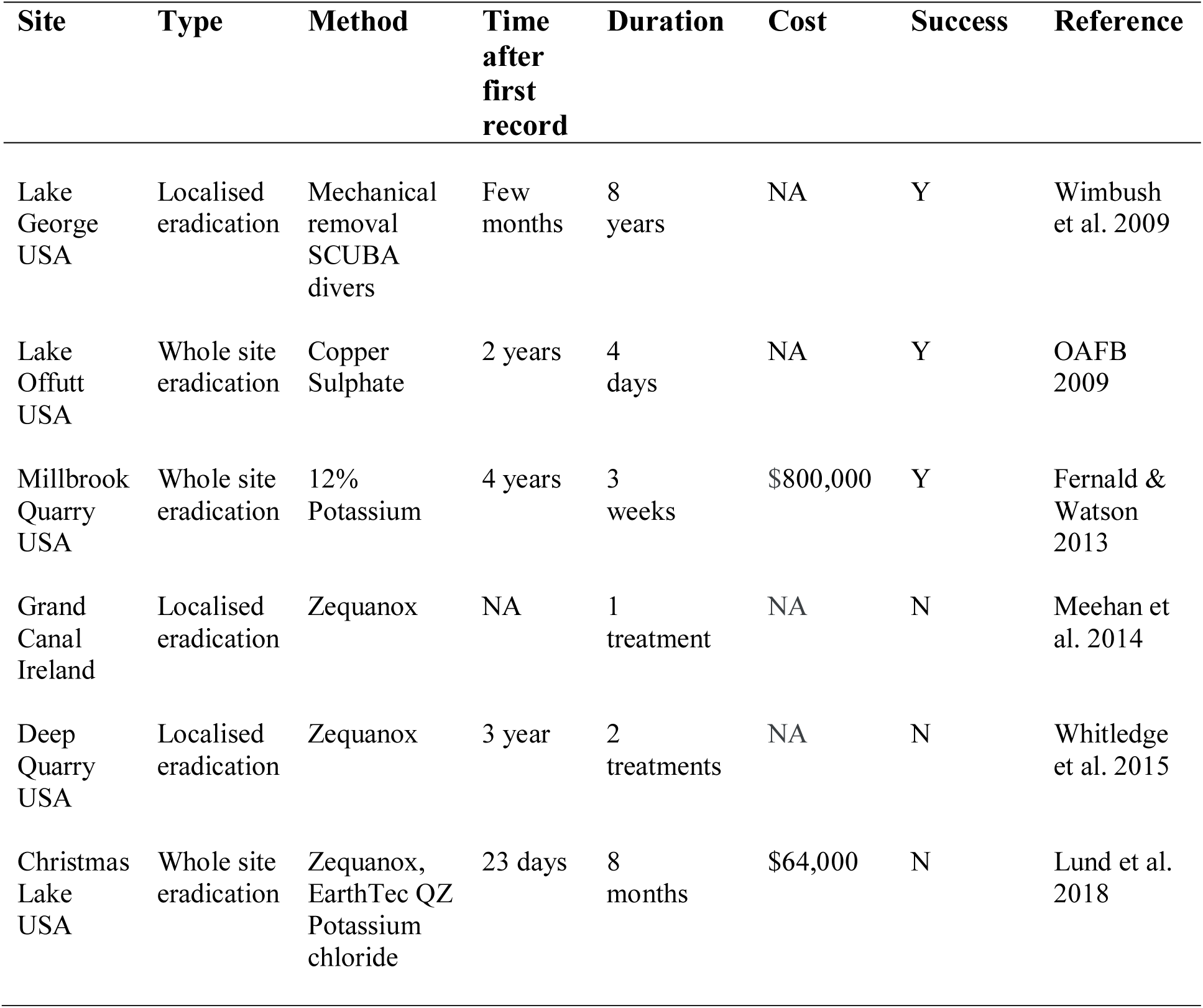
Examples of eradication programmes of zebra mussel.

| Site | Type | Method | Time<br>after<br>first<br>record | Duration | Cost | Success | Reference |
| --- | --- | --- | --- | --- | --- | --- | --- |
| Lake George<br>USA | Localised eradication | Mechanical removal<br>SCUBA divers | Few months | 8 years | NA | Y | Wimbush et al. 2009 |
| Lake Offutt<br>USA | Whole site eradication | Copper Sulphate | 2 years | 4 days | NA | Y | OAFB 2009 |
| Millbrook Quarry<br>USA | Whole site eradication | 12% Potassium | 4 years | 3 weeks | \$800,000 | Y | Fernald & Watson 2013 |
| Grand Canal<br>Ireland | Localised eradication | Zequanox | NA | 1 treatment | NA | N | Meehan et al. 2014 |
| Deep Quarry<br>USA | Localised eradication | Zequanox | 3 year | 2 treatments | NA | N | Whitledge et al. 2015 |
| Christmas Lake<br>USA | Whole site eradication | Zequanox, EarthTec QZ<br>Potassium chloride | 23 days | 8 months | \$64,000 | N | Lund et al. 2018 |

**Table 1.** Estimated cohort composition of successive waves of zebra mussels colonising experimental panels in Cardiff Bay over a 12 month period, based on mixture analysis of shell length frequency data (n = 2,785).

| Cohort | Approx. Age<br>(days) | Mean size<br>(mm) | Size<br>range (mm) | SD | % Frequency |
| --- | --- | --- | --- | --- | --- |
| 1 | 66 | 12.2 | 11-12 | 0.85 | 7.3 |
| 2 | 132 | 15.4 | 9-17 | 2.02 | 47.2 |
| 3 | 198 | 18.6 | 18-19 | 0.57 | 17.3 |
| 4 | 264 | 21.5 | 20-22 | 0.84 | 12.2 |
| 5 | 330 | 25.0 | 23-31 | 2.12 | 15.9 |

Several traits of the zebra mussel make their eradication particularly challenging: (1) the species is highly fecund (up to 1 million eggs per female), (2) has a planktonic veliger stage which can travel over great distances and survive for weeks (Minchin et al. 2002), (3) displays high tolerance to a wide range of temperatures (Spidle et al. 1995), salinities (Kilgour et al. 1994) and pH values (Bowman & Bailey 1998), and (4) it has a tendency to aggregate in enormous beds (up to 32,500 individuals/m^2^ -Berkman et al. (1998)) on different types of substrates. Zebra mussel invasions have been facilitated by many anthropogenic actions, including the building of canals and channels that connect formerly isolated water bodies (Decksbach 1935), boat trading (Kearney & Morton 1970) and aquatic leisure activities (Kinzelbach 1992). These can make eradication particularly difficult, as the risk of recolonization from connected water ways is always high (Mari et al. 2011). In Britain, the first record of zebra mussel dates back to 1824, but it is only since 2000 that the species has started to spread rapidly and cause widespread ecological damage, a pattern that cannot be explained simply by increasing public awareness (Aldridge et al. 2004) and that remains unclear.

The recruitment and demography of zebra mussel have been well studied in North America (Chase & Bailey 1999; Martel et al. 1994), but there is relatively little information on the dynamics of the species in recently invaded waters in Europe (Alix et al. 2016; MacNeil et al. 2010), nor is it clear how populations are structured during the initial stages of the invasion, when boom and bust dynamics have been reported (Strayer et al. 2017).

One pressing issue with the control of zebra mussel is to assess to what extent incomplete eradication (i.e. partial removal) is useful in controlling population growth and limit dispersal, or on the contrary, may cause more harm than good if populations simply bounce back in greater numbers. Under controlled environments, mitigation measures can help reduce the abundance of zebra mussel (Luoma et al. 2018; Waller & Bartsch 2018), which could help reduce impacts (Fernald & Watson 2013; Wimbush et al. 2009). However, information on natural systems is very scant and models have typically low predictive power to predict zebra mussel dispersal patterns (Rodriguez-Rey et al. under review). The ability of zebra mussel to recover from partial removal will likely depend on seasonal patterns of growth and recruitment, which have been correlated with seasonal temperatures (Allen et al. 1999) and (Churchill et al. 2017), chlorophyll-a (Churchill et al. 2017), but also with calcium, alkalinity, and total hardness (Hincks & Mackie 1997). Mortality and recruitment appear to be influenced by fluctuations in temperature, but also by population size structure (Allen et al. 1999), and there is some evidence that settlement of new juveniles is negatively affected by the density of established adult mussels (Nalepa et al. 1995), suggesting the existence of negative density-dependence processes.

The recruitment and demography of zebra mussel have been well studied in North America (Chase & Bailey 1999; Martel 1993; Martel et al. 1994; Nalepa et al. 1993) but there is relatively little information on the dynamics of the species in recently invaded waters in Europe (Alix et al. 2016; MacNeil et al. 2010). Nor is it clear how populations are structured during the initial stages of the invasion, when boom and bust dynamics might be expected (Strayer et al. 2017), and where a better understanding of growth and recruitment could make control measures more efficient.

### Aims and objectives

We monitored the colonization and growth of zebra mussel in experimental panels submerged at different depths in Cardiff Bay (Wales, UK), an amenity lake where the species is spreading and causing increasing damage. Zebra mussels were first recorded in Cardiff Bay in 2004 (although they may have been introduced a year earlier), and have spread rapidly ever since, being now present throughout the Bay (Alix 2010; Alix et al. 2016; Wood et al. 2015). The objectives of the study were three (1) to assess the extent of seasonal and spatial variation in the growth and settlement rates of zebra mussel in a recently colonized artificial lake area, (2) to identify the conditions that are most favourable for zebra mussel production, and (3) to test for the existence of density-dependent growth. Ultimately, it was hoped that the results of this study might serve to inform the development of more efficient control measures by targeting those periods and locations where zebra mussel production is highest.

## Material and Methods

### Study area

The study site, Cardiff Bay, is a 2.0 km^2^ amenity lake (depth = 4-7 m) located in Cardiff (Wales, UK) and fed by two rivers (River Taff and River Ely). It was built between 1994 and 1999 as part of a regeneration project of the old docklands areas of Cardiff and Penarth. The site has been described in detail by (Alix 2010; Alix et al. 2016).

### Sampling strategy

We deployed five experimental buoys in different parts of Cardiff Bay (Figure 1.), each buoy consisting of a weighted rope and three white plastic panels (A4 size, 210 × 297 mm) set at the surface (0m), 1m and 3m depth (Figure 2.). The buoys were deployed on June 2017 and were monitored monthly until May 2018.

**Figure 1.**
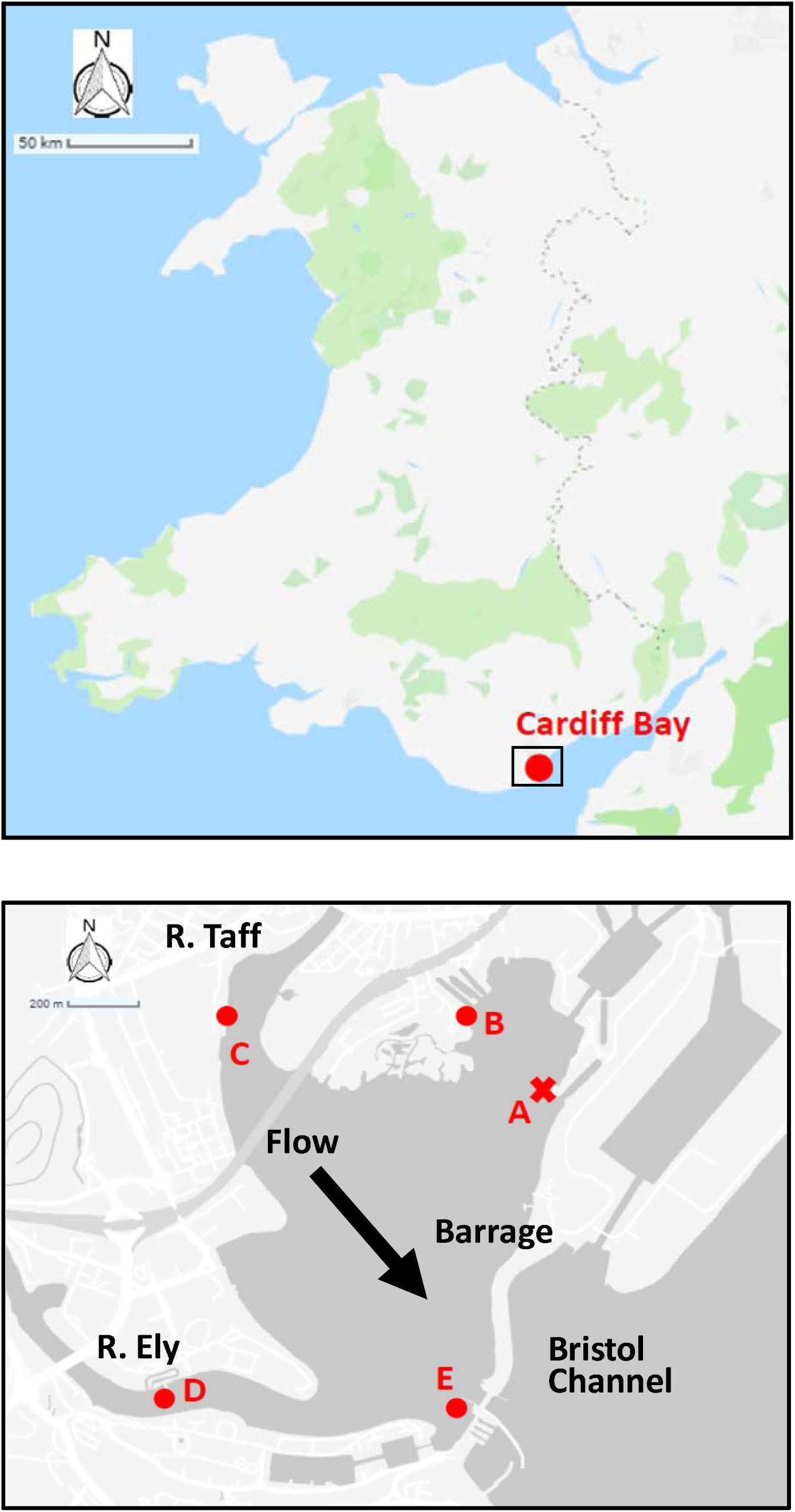
Study site for zebra mussel population dynamics (Cardiff Bay, UK) showing location of experimental buoys (A-E) used to sample new recruits. Buoy A was lost during the first month and was excluded from analysis.

**Figure 2.**
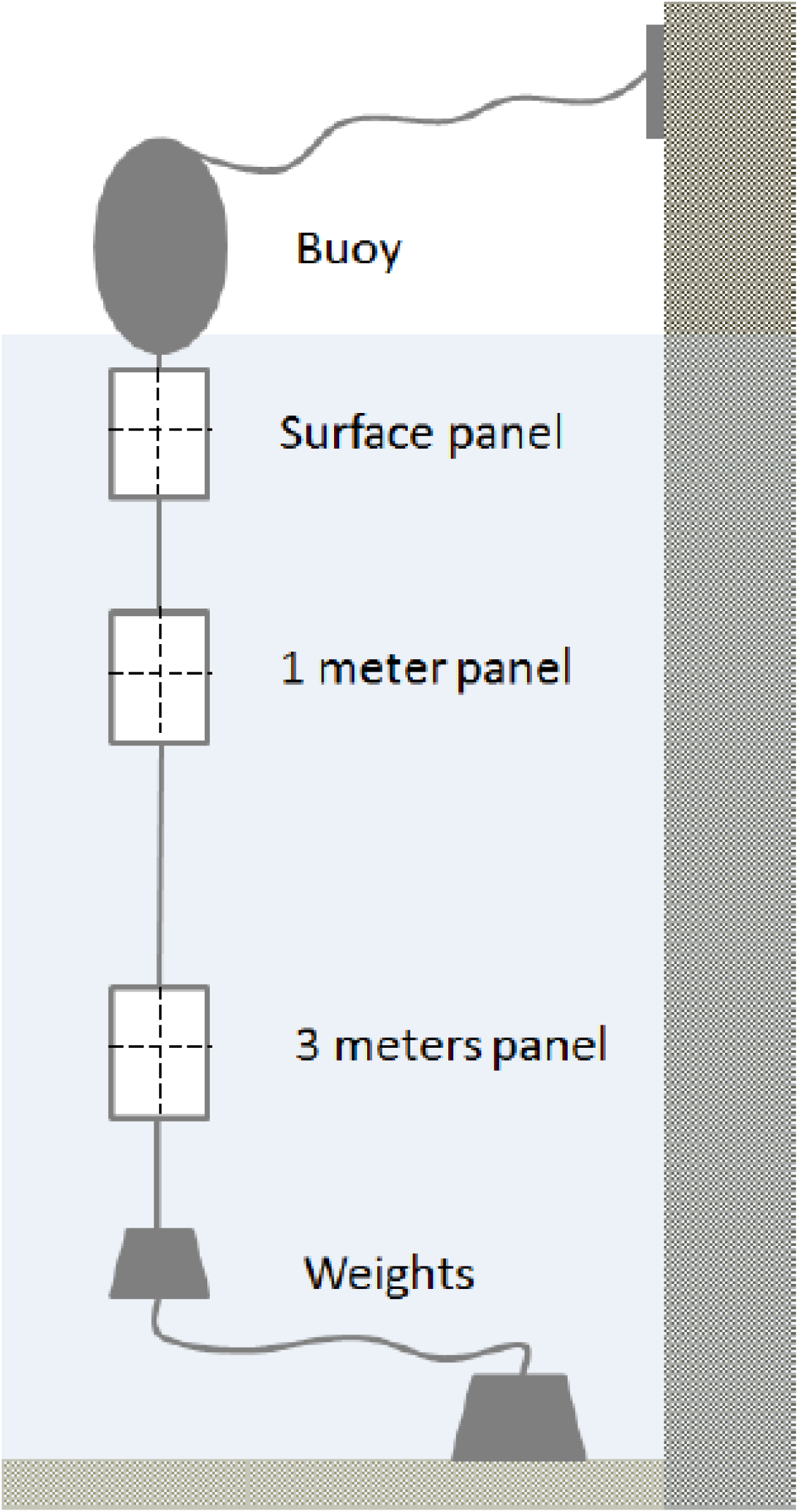
Experimental buoys used to assess the influence of water depth and density on zebra mussel population dynamics. Each month 25% of the area of each panel (156 cm^2^) was scrapped clean and all the attached zebra mussels were counted and measured.

An area corresponding to 25% of each panel (i.e. 156 cm^2^) was scraped clean every month and all attached mussels were counted and measured. These are referred to as ‘clean samples’ and provide data on the number and size of new recruits. A total of 141 scrape samples were collected in this way over the 12 months of the study, 43 of which contained zebra mussels (30.5%). In addition, 81 scrape samples were obtained from colonised sections of the panel (i.e. let undisturbed, never scraped before) 5 months after the start of the study; each month a different area of the panel was scraped, 67 of which contained zebra mussel (82.7%). Specimens were preserved in 70% ethanol and brought to the laboratory where they were counted and measured (shell length along the longest axis, mm).

During the monthly monitoring, water parameters (temperature, DO concentration, DO %, salinity, conductivity, pH, turbidity) were measured at 0m, 1m and 3m depth at each sampling station with a probe (YSI Water Quality Sonde,6600 EDS V2, USA). Due to bad weather, buoy A was lost during the first month and was excluded from analysis, and buoy B was lost during the last month of the experiment but data were available for 11 of the 12 months of the study.

### Statistical Analysis

Statistical analyses were carried out using R version 3.3 (Core Team, 2017) and PAST v. 3.2.2 (Hammer et al. 2001). We used mixture analysis on shell length at the end of the growing season to estimate the number of different cohorts (age classes) that had colonised our experimental panels. For this, we varied the number of putative cohorts from 1 to 8 and chose the most likely number based on changes in AIC values (Hammer et al. 2001). We used linear models to examine variation in mussel size and density using month, depth and site as predictors. To examine the influence of water parameters, we used principal component analysis using the *prcomp* function in the factoextra R package (Kassambara & Mundt 2017) and used the coordinates of the first principal component as predictors of mussel size and density in a linear mixed effect model using sampling station as a random factor, as above. Model simplification was achieved by examining changes in AIC using the *step* and *dredge* functions, followed by Maximum Likelihood comparisons of nested models with the *anova* command.

To test for evidence of density-dependence, we tested if density was a significant predictor of the average size of mussels in each sample, taking into account the effects of season, water depth and variation among sites. We carried out an analysis separately for one month old mussels (originating from our monthly scrape panels) and for mussels sampled from undisturbed panels at the end of the growing season. As the relation between density and size was not linear, we employed generalised additive modelling (GAM) using a penalized regression spline fitted by REML in the *mgcv* package to account for non-linearity (Wood 2001), and dropped non-significant terms from the final model. We excluded site B from analysis as there was no colonization of surface panels, and used the *gam.check* command to assess departures from model assumptions.

## Results

### Variation in water chemistry

Water chemistry changed both seasonally (Figure 3.) and spatially across Cardiff Bay. Water temperatures peaked in Jul-September (max = 20.4 C) and reached a low in March (min = 5.6 C; month *F*_11,207_ = 1055.6, *P* <0.001), being generally warmest at the mouth of the River Ely (site D) and the barrage (site E), and coldest at the mouth of the River Taff (site C) and the inner harbour (site B, *F*_3,207_ = 23.1, *P* <0.001). Dissolved oxygen reached a minimum in July (min = 6.1 mg/L), coinciding with the warmest temperature (*F*_11,207_ = 1055.6, *P* <0.001), and was highest at the outlet of the Bay (site E), and lowest at site D (*F*_3,207_ = 23.1, *P* <0.001). The four other water chemistry parameters also varied significantly from month to month, as well as from site to site (conductivity: month *F*_11,207_ = 1055.6, *P* <0.001, site *F*_3,207_ = 23.1, *P* <0.001; pH: month *F*_11,207_ = 1055.6, *P* <0.001, site *F*_3,207_ = 23.1, *P* <0.001; salinity: month *F*_11,207_ = 1055.6, *P* <0.001, site *F*_3,207_ = 23.1, *P* <0.001; turbidity: month *F*_11,207_ = 1055.6, *P* <0.001, site *F*_3,207_ = 23.1, *P* <0.001). However, no significant variation in water chemistry was found with respect to water depth, at least within the first 3 metres (P > 0.5 in all models).

**Figure 3.**
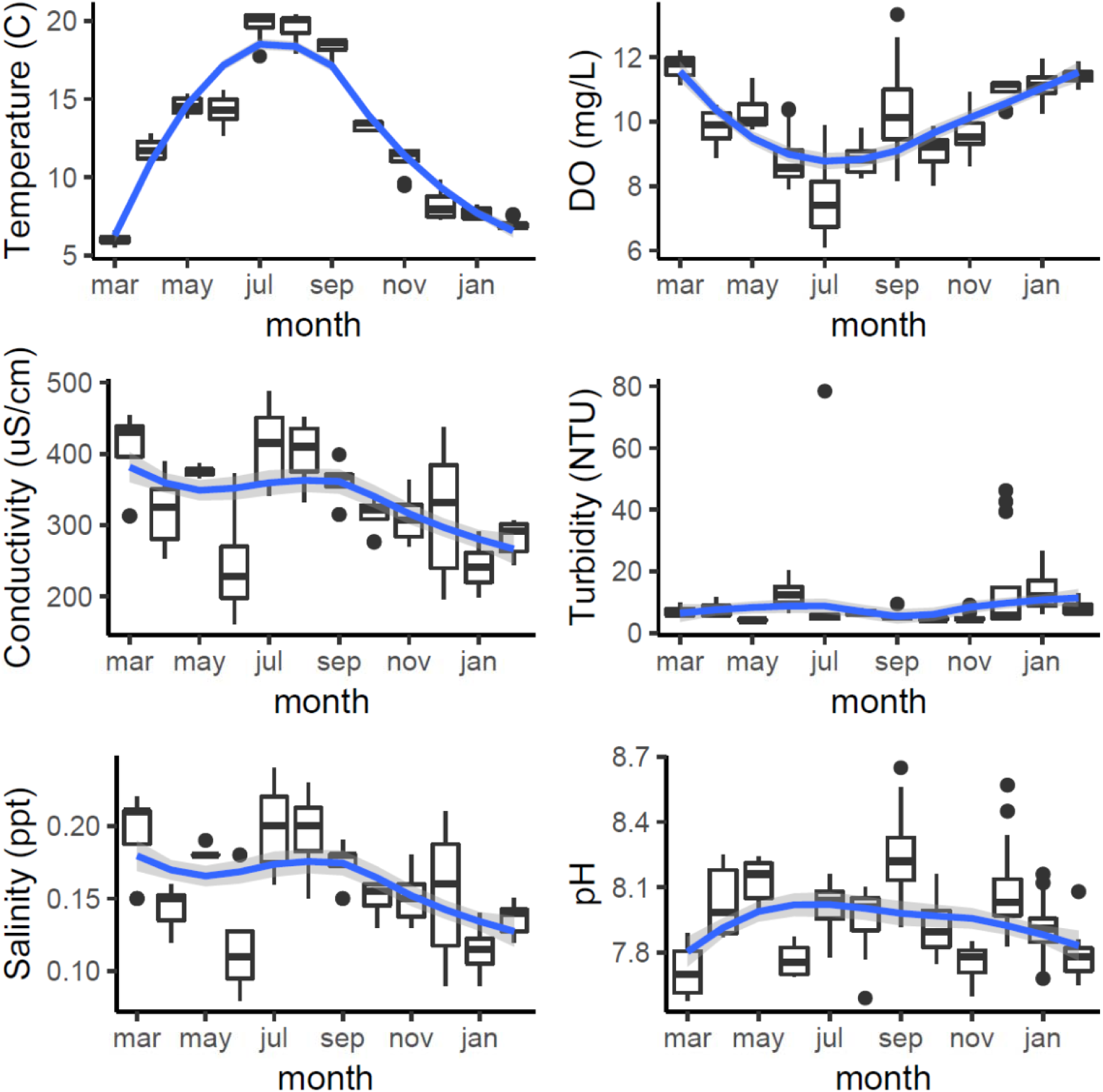
Monthly variation in water chemistry in Cardiff Bay during the course of the study showing median values and interquartile range (boxplots) and smoothed fitted trends with 95CI envelopes (blue and grey bands).

Principal component analysis indicated that the first component (PC1) accounted for 39% of the variation in water chemistry parameters, but was not a significant predictor of either the average size (*t* = 0.787, *P* = 0.434) or density (*t* =2.46, *P* = 0.750) of new recruits colonising the experimental panels.

### Density and settling rates

Densities of one month old mussels sequentially sampled during the reproductive season varied between 0 in June and 1.8 individuals/cm^2^ in September (Figure 4.). Colonization of the experimental panels began in July, peaked during September and October, and then decreased rapidly, so that by November no new recruits were detected in any of the panels (Figure 4.). Densities of one month old zebra mussel varied significantly between months (*F*_1,134_ = 15.8, *P*<0.001), sampling sites (*F*_3,134_ = 4.02, *P*=0.009), and depths (*F*_2,134_ = 3.46, *P=*0.03). The surface panels had the lowest number of recruits, while the deepest panel had the highest.

**Figure 4.**
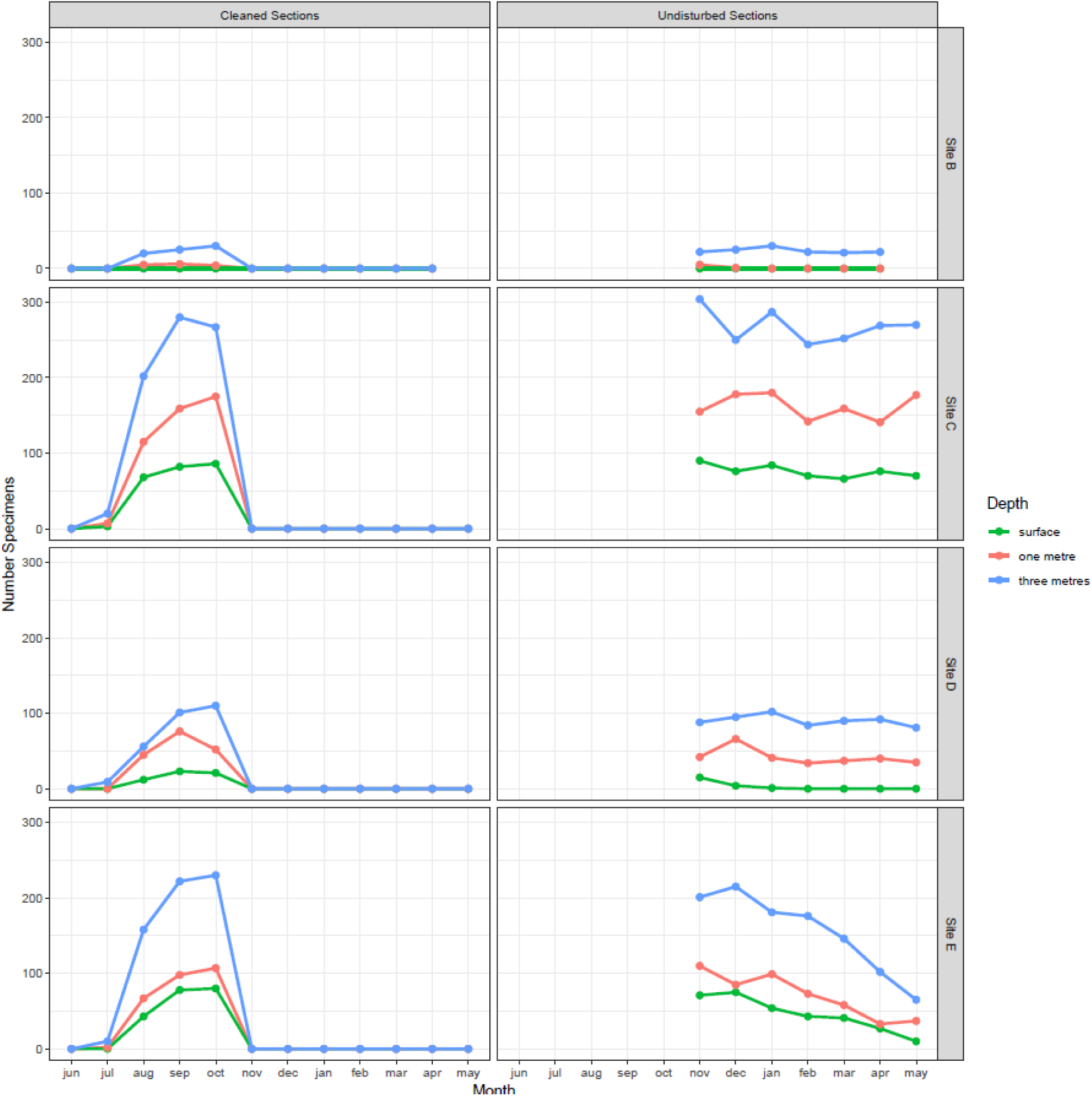
Monthly variation in the number of zebra mussel found at different depths and at different locations in Cardiff Bay. Left panel shows results for panels scrapped clean every month during the growing season and right panel the results for panels left undisturbed and sampled after the end of the growing season.

Densities in the undisturbed panels, sampled over the winter, also varied between sampling sites (*F*_3,75_ = 87.3, *P*<0.001) and depths (*F*_2,75_ = 71.2, *P*<0.001), but were stable across time (month *F*_6,69_ = 1.67, *P* = 0.142), confirming the lack of recruitment observed between November and May. In general, the highest densities and settling rates were found at the mouth of the River Taff (site C) and at the outlet at the barrage (site E), while the lowest were found at the inner harbour (site B).

Settling rates across the Bay followed a marked seasonal cycle (Figure 5.), closely tracking variation in water temperature, beginning in June when temperature reached 14 C, peaking in August and September, and then ceasing when temperature dropped below 14 C in October-November. Across sampling stations, settling rates were 0.42-0.62 indiv cm^2^ month^−1^, with peaks of 1.790 indiv cm^2^ month^−1^. This is equivalent to 4,200-6,200 mussels per m^2^ (peaks of 18,000 indiv/m^2^)

**Figure 5.**
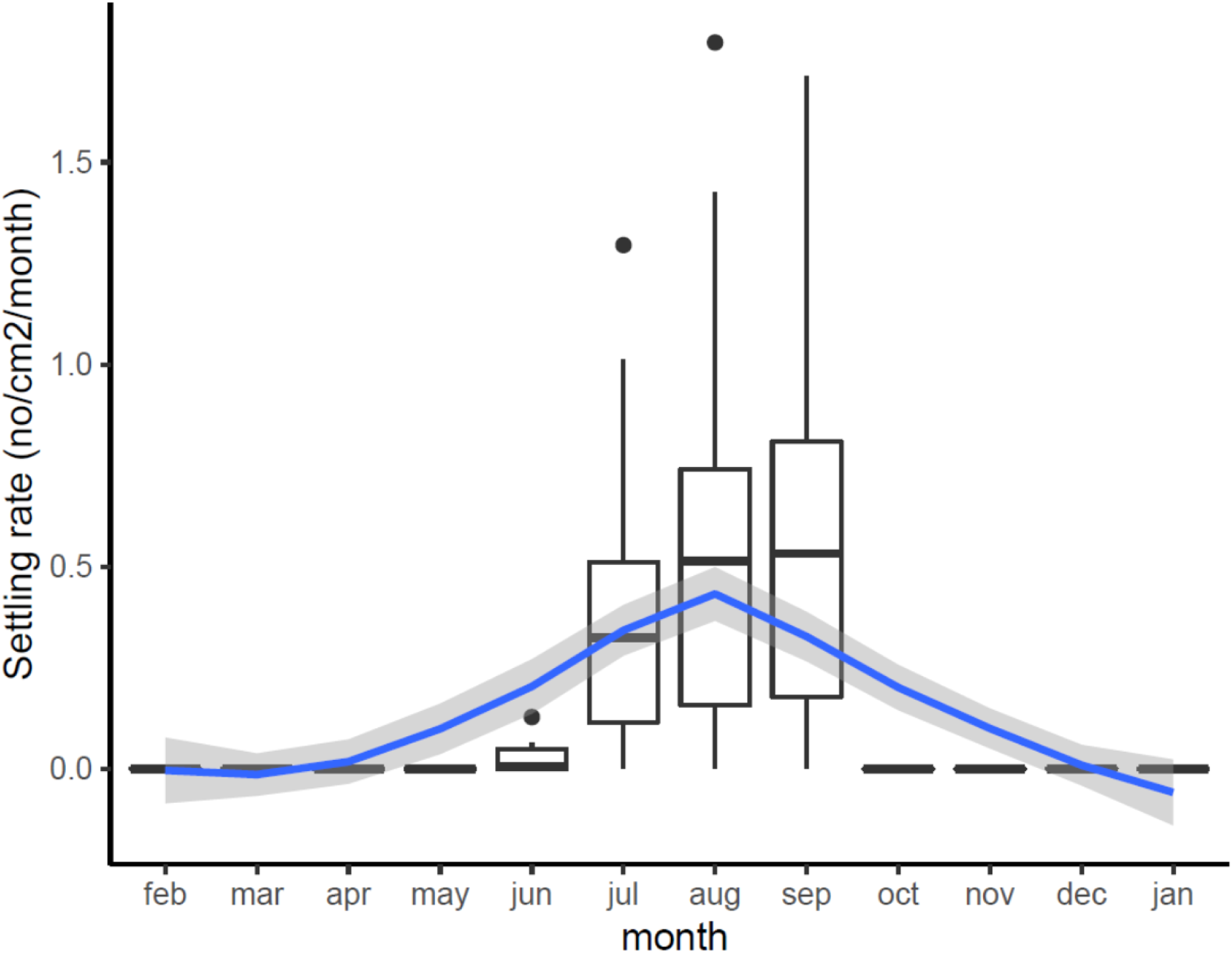
Estimated variation in monthly settling rates (No. new recruits/cm^2^/month) across Cardiff Bay showing median values and interquartile range (boxplots) and smoothed fitted trend with 95CI envelope (blue and grey bands).

### Determinants of mussel size and growth

The average size of one month old mussels colonizing the clean panels during the reproductive season varied between 9mm in July to 21mm in October, and differed significantly between months (*F*_3,1377_= 700.7, *P*<0.001), sampling sites (*F*_3,1377_ = 293.3, *P*<0.001), and also with depth (*F*_1,1377_ = 385.4, *P*<0.001; Figure 6.). The largest mussels were found at the mouth of the River Taff (site C) and at the outlet at the barrage (site E), while the smallest ones were found at the mouth of the River Ely (site D). Growth increased rapidly from July to October, and then plateaued for the rest of the year. The size of mussels was largest at 3 metres (95 CI = 15.7-16.3 mm) and smallest at the surface (95CI = 14.0-14.8 mm). Such variation persisted in the undisturbed panels over the winter, after the reproductive season, as mussel size continued to vary significantly between months (*F*_6,2774_ = 11.6, *P*<0.001), sampling sites (*F*_3,2774_ = 170.9, *P*<0.001) and also with depth (*F*_1,2774_ = 206.8, *P*<0.001; Figure 6.). Thus, the average size of mussels in Apr 2018, 10 months after the buoys were first deployed, was still significantly smaller at the surface (mean = 17.0mm) than at 1m depth (mean = 18.3mm) and at 3m depth (mean = 18.4 mm; Tukey HSD *P* adj = 0.003), which were not different among themselves (Tukey HSD, *P* adj = 0.732).

**Figure 6.**
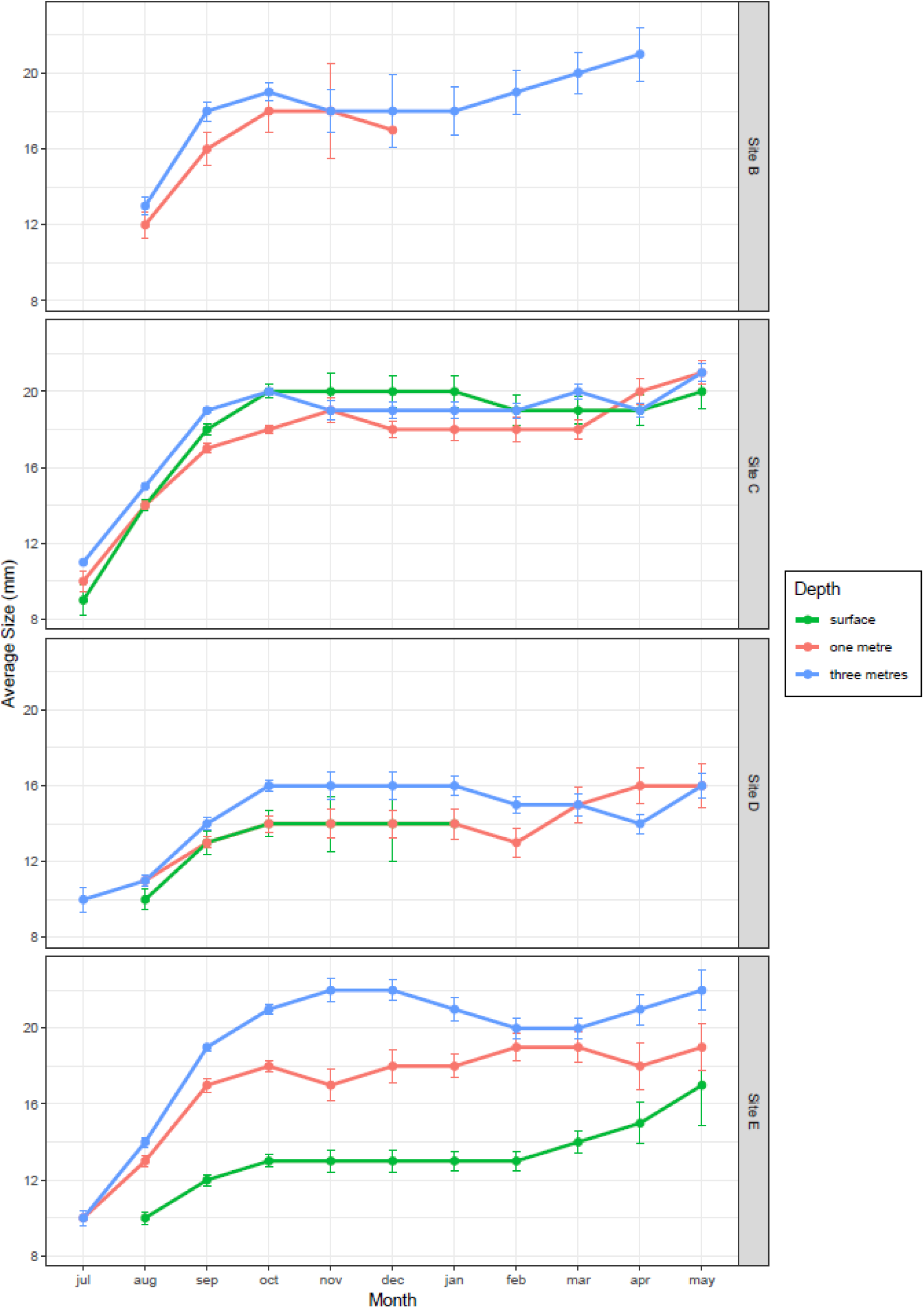
Growth trajectories of zebra mussel at different depths and locations in Cardiff Bay (mean ± 95CI).

### Cohort analysis

Inspection of experimental panels revealed that new recruits were only found during July to October, suggesting that the reproductive season in Cardiff Bay likely extended from May or June to September. Results from mixture analysis suggest that the most plausible number of discrete cohorts colonising the experimental panels over the course of the study was 5 age classes (Figure S1), with an estimated age of approximately 2 months for the youngest settlers (size 11-12 mmm) to 330 days for the oldest ones when zebra mussels had already attained a size of 23-31 mm (Table 2.). The distribution of cohorts varied significantly among sites (Chi-squared = 387.5, df = 12, *P* < 0.001) and there were comparatively more younger settlers at the warmest sites (sites D and E) than at the coldest ones (sites C and B, Figure 7.).

**Figure 7.**
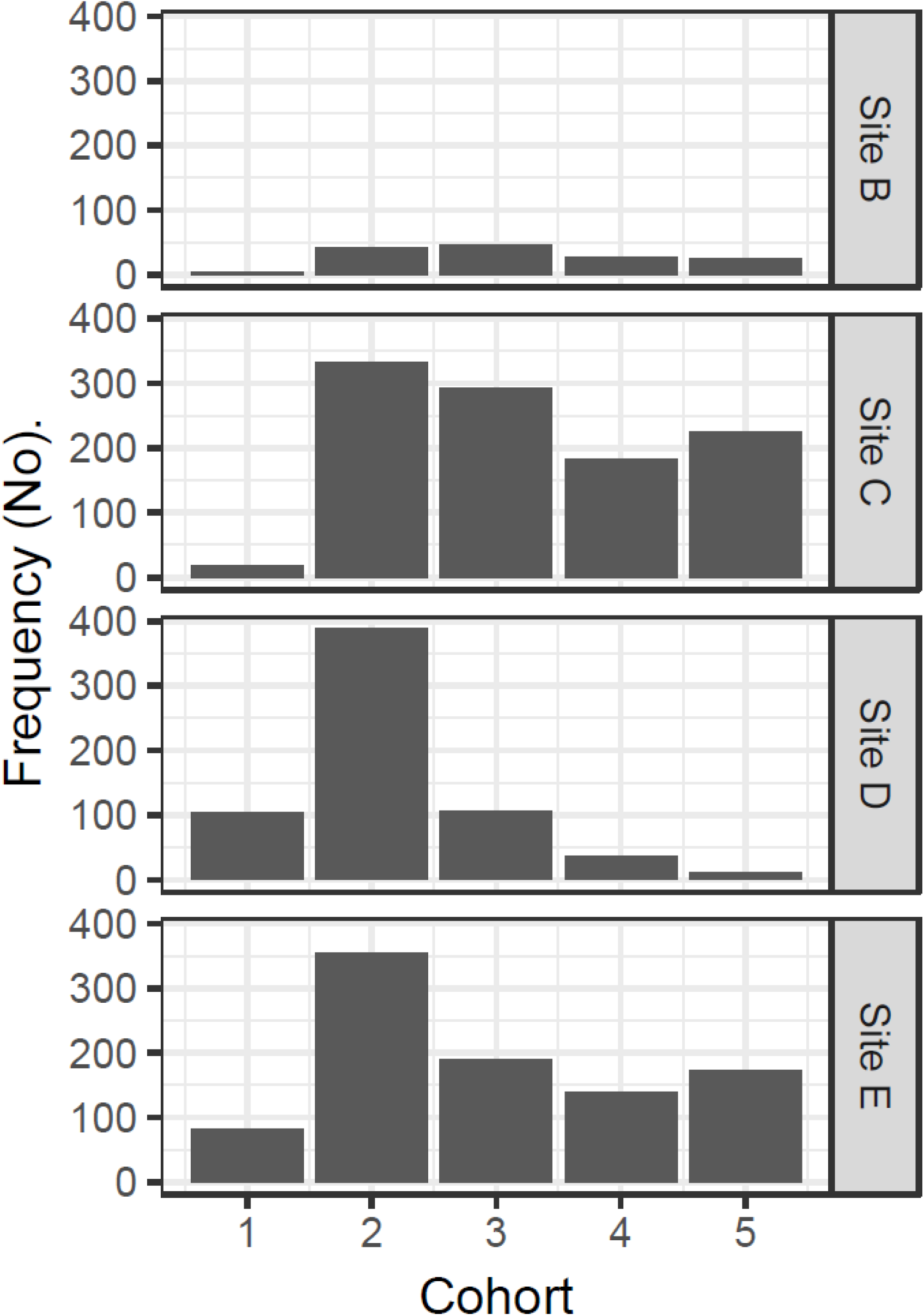
Site variation in the relative frequency of different cohorts colonizing the experimental panels in Cardiff Bay derived from mixture analysis.

### Density dependence

The size of one month old mussels (i.e. new settlers successively sampled from cleaned panels) was not affected by density, once the effects of site, water depth and month of sampling had been statistically controlled for (density *F*_1,29_ = 0.427, *P* = 0.519). However, density was a significant predictor of mussel growth in the undisturbed panels (GAM estimates for smooth terms; density, *F*_3.357,4.090_ = 4.433, *P* = 0.004; density x site C, *F*_5.405,6.099_ = 14.881, *P* < 0.001; density x site D, *F*_3.154,3.606_ = 16.688, *P* <0.001; density x site E, *F*_1,1_ = 3.732, *P* =0.06; parametric terms, depth 1m estimate = 0.466, SE= 0.05, *t* = 9.266, *P* <0.001; depth 3m estimate = 0.759, SE = 0.07, *t* = 10.267, *P* <0.001). The model explained 91.6% of deviance in mussel size, of which 39.4% was explained by density alone. Across sites, mussel size increased with density (Figure 8.), but there were significant differences between sites. Thus, at sites with high recruitment (site C, mouth of River Taff; site E, outlet of the barrage, Figure 5.) mussel size decreased at high densities, whereas at the site with low recruitment (site D, mouth of River Ely) the opposite was found (Figure 5.). This suggests the existence of positive density-dependent growth at low densities, and negative density-dependent growth at high densities.

**Figure 8.**
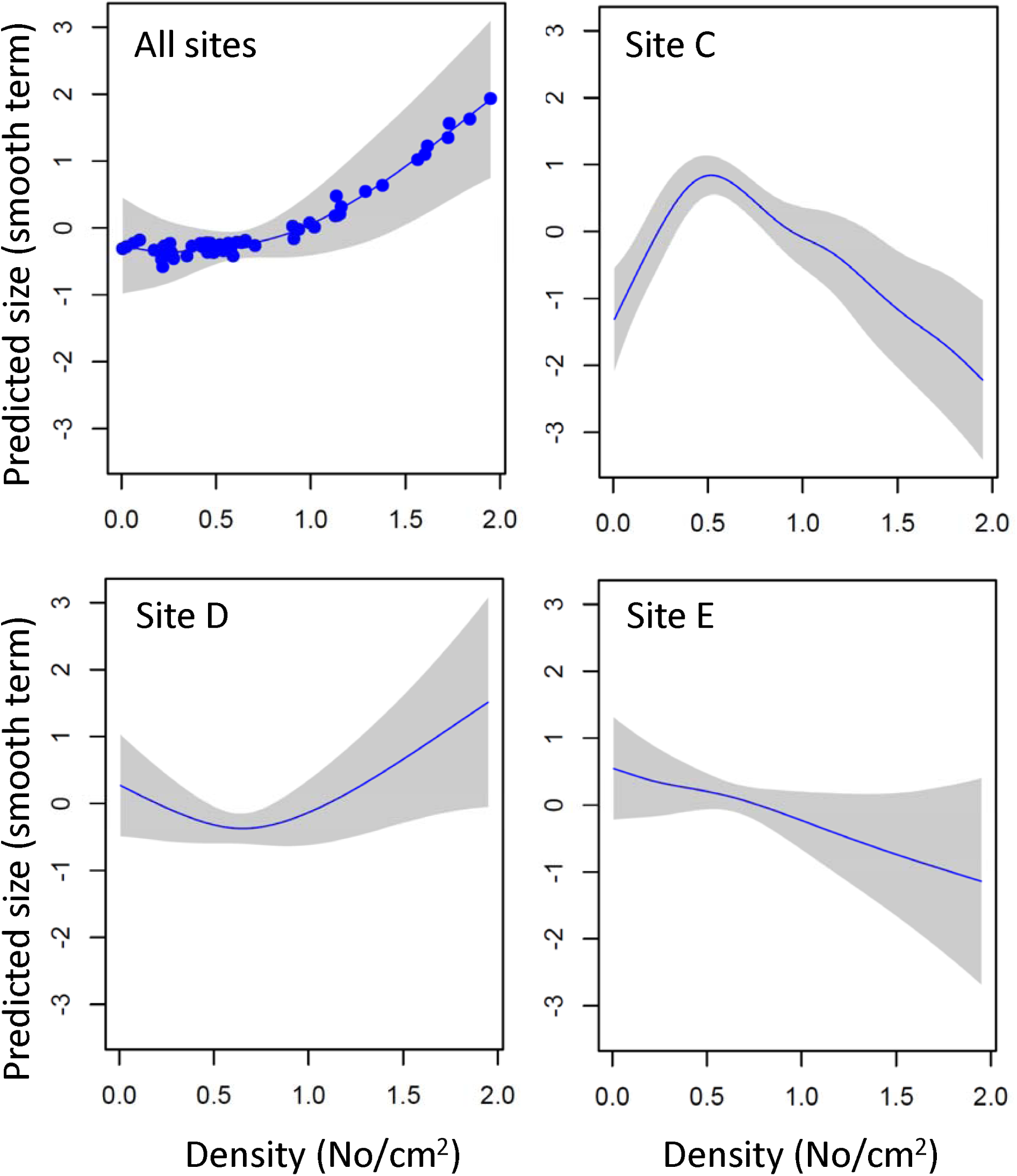
Density-dependent changes in the growth of zebra mussel in different parts of Cardiff Bay. Shown are predicted sizes (GAM smoothed term centered around zero) and 95 CI envelopes across all sites, at sites with high recruitment (Sites C and E) and at a site of low recruitment (site D).

## Discussion

Our study indicates that within 14 years of being invaded, Cardiff Bay has a large, established population of zebra mussel, confirming the conclusions of a previous survey of veliger density using a different sampling approach (Alix et al. 2016). Our study also suggests that zebra mussel may be spatially structured into different cohorts that grow and settle at different rates. This has implications for management because if density, settling rates, and growth vary spatially among locations, this creates the conditions necessary for zebra mussel metapopulations to evolve, which would make control measures considerably more challenging (Mari et al. 2014).

By carrying out monthly scrapes in artificial substrates, we have shown that zebra mussel begin to settle in July, one month after we deployed the experimental panels, and continued until October, with no evidence of new colonisations from November to May. Density and growth followed the same pattern, increasing over the summer and ceasing in October, after which no further recruitment or growth occurred. These findings are similar to those of previous studies (Alix et al. 2016; Fong et al. 1995; Ram et al. 1996). The fastest growth was generally observed at the deepest waters (3 meters), which also coincides with the highest settling rates of new recruits. This strongly suggests that conditions that favour growth of zebra mussel also favour their survival. However, no significant differences in water chemistry were found within the first 3 metres, despite a large variation in growth and settling rates within the water column and our index of water chemistry (PC1) did not explain the size or abundance of mussels, despite large variation in water parameters among sites. This suggests that factors other than water chemistry control growth and colonisation of zebra mussel, most likely physical disturbance, predation pressure, and food abundance. Water parameters for Cardiff Bay are within the optimal values for zebra mussel (Bowman & Bailey 1998; Kilgour et al. 1994; Spidle et al. 1995). However, the Bay is fitted with a bottom aeration system to maintain high dissolved oxygen and permit the passage of migratory Atlantic salmon and brown trout (Alix et al. 2016), and as a result, water is more mixed than would normally be, which may explain the apparent lack of stratification in water parameters (Alix 2010).

The low settling rate of mussels found in surface samples has been noted previously in laboratory and field studies (Alix et al. 2016; Kobak 2001; Kobak 2004). Velgers appear to be absent from the upper 50 cm of the water column in Cardiff Bay (Alix et al. 2016). Avoidance of surface waters appears to be related to light intensity (Kobak 2001; Seaver et al. 2009) and may confer mussels some protection against bird predators and desiccation caused by fluctuating water levels. For example, Alix (2010) reported that waterfowl fed on surface mussels in Cardiff Bay and has also been found that waves reduce settlement rates (Chase & Bailey 1999; Kobak 2004), which may explain the low abundance of mussels in our surface panels. Mean settling rates varied between 4,200 and 6,200 mussels m^−2^ month^−1^, which are similar to those reported for well established populations elsewhere (Cleven & Frenzel 1993; Mackie & Schloesser 1996; Stanczykowska & Lewandowski 1993), and are also consistent with adult densities of 450–5,100 mussels m^−2^ estimated at Cardiff Bay during 2006-2009 (Alix 2010; Alix et al. 2016). However, the peak of 17,960 adult mussels m^−2^ month^−1^ recorded on September 2017 at a depth of 3m at the mouth of the river Taff (site C) is over 2.3 times higher than the highest value reported previously for Cardiff Bay (7,700 indiv. m^−2^ -Alix et al. (2016)), and also higher than the highest density ever recorded in Britain (11,000 ind. m^−2^, -Aldridge et al. (2004)). This may indicate that the zebra mussel population in Cardiff Bay is increasing despite the removal of approximately 4 tonnes of mussels every year (Alix et al. 2016).

Our results suggest the existence of five distinct cohorts resulting from the same spawning season, that our data suggest it extends from June to September (Figure 5.). This is in agreement with results from previous surveys on veliger density that also indicated a reproductive season extending from late May/June to late September/October for Cardiff Bay (Alix et al. 2016). The presence of multiple cohorts from the same spawning season has not been reported previously but is consistent with results from the laboratory that indicate that the release of gametes occurs over 2–6 spaced events (Haag & Garton 1992; Walz 1978). It can also result from spatial variation in the timing of reproduction and growth across the Bay, as well as from dispersal at the post-settlement adult stage. Dispersal of zebra mussels is mostly through the planktonic veliger stage but settled adults can also disperse. Adults may choose to dislodge when conditions become unsuitable and be transported long distances attached to macrophytes and other vectors, and also drift using the byssus to gain buoyancy (Kobak 2001; Martel 1993). This means that colonization is not restricted to the veliger stage immediately after reproduction, but that it can be extended beyond this phase. Post-settlement dispersal, therefore, coupled with variation in timing of reproduction among locations and multiple releases of gamete could give rise to multiple cohorts and different waves of settlers, as observed in our study. More generally, variation in habitat quality, timing of reproduction and demography creates the conditions necessary for source and sink metapopulations to develop (Vanhaecke et al. 2012) which may need to be taken into account in the management of zebra mussel. For example, our results suggest that different density dependent mechanisms may operate in different habitats, once the effects of depth and month of sampling had been taken into account.

In general, mussel size increased with density, but there were marked differences between sites. There was positive density-dependent growth at sites with low densities, and negative density-dependent growth at sites with high densities, suggesting some form of population regulation, presumably caused by competition. Zebra mussel settling rates have been reported to be lower on substrate densely populated by adults (Nalepa et al. 1995) which is consistent with negative density dependence. This can have implications for management because removal could be effective or counter-productive depending on conditions. The cost of controlling zebra mussel populations is generally high (Adams & Lee 2012) and while different eradication methods have been tested in the laboratory (Claudi et al. 2014; Costa et al. 2011; Luoma et al. 2015; Watters et al. 2013) these are not always successful in the field (Table 1.). The development of more efficient eradication methods should benefit from insights into natural factors regulating population growth (Lund et al. 2018). Currently, control measures against zebra mussel are limited to the use of various chemicals (Glomski 2015), ultraviolet radiation (Lewis & Whitby 1997) and mechanical removal and drying (Durán et al. 2010). These aim to reduce populations size, but our results may help explain why removal may not always work. Zebra mussels often follow boom and bust population dynamics (Casagrandi et al. 2007; Strayer et al. 2017), which has previously been associated with size selective predation by birds (Pedroli 1977; Wisniewski 1974). We suggest that when eradication is not possible, local zebra mussel populations dynamics should be considered before embarking on partial removal that may prove expensive, ineffectual, and may in some cases enhance production and aggravate the problem

In conclusion, our results indicate that the zebra mussel is a well established aquatic invader in Cardiff Bay and despite periodic removal, its numbers appear to be growing as evidenced by having the highest densities recorded in Britain to date, and also higher than previous estimates for this artificial water body. Given the overriding effect of water temperature on reproduction, under current predictions of climate change the spawning period of zebra mussel in Cardiff Bay will likely extend, which may result in even higher production. This makes the search for more effective control measures paramount. Based on the observed seasonal pattern of growth and recruitment, we suggest that control measures might benefit from targeting the summer and early autumn for removal of zebra mussel, as this appears to be the most critical period for the colonisation and dispersal of this invasive species. We also suggest that control actions should target suitable structures located 3 m deep or deeper, as this appears to be the zone where most of the zebra mussel production occurs. However, the existence of contrasting patterns of density dependent growth (positive at sites of low recruitment and negative at sites of high recruitment) suggests that removal of adult mussels may help curtail biomass production at some sites but may enhance it at other sites. The existence of significant spatial variation in growth and settling rates across the Bay, coupled with multiple cohorts, suggest that zebra mussel might be structured as a metapopulation governed by source and sink dynamics. Although our study cannot resolve this possibility, this merits further attention and could be addressed by using molecular markers to determine patterns of gene flow, as shown for the larval dispersal of other aquatic species (e.g. Vanhaecke et al. (2012)).

---

**Figure S1.**
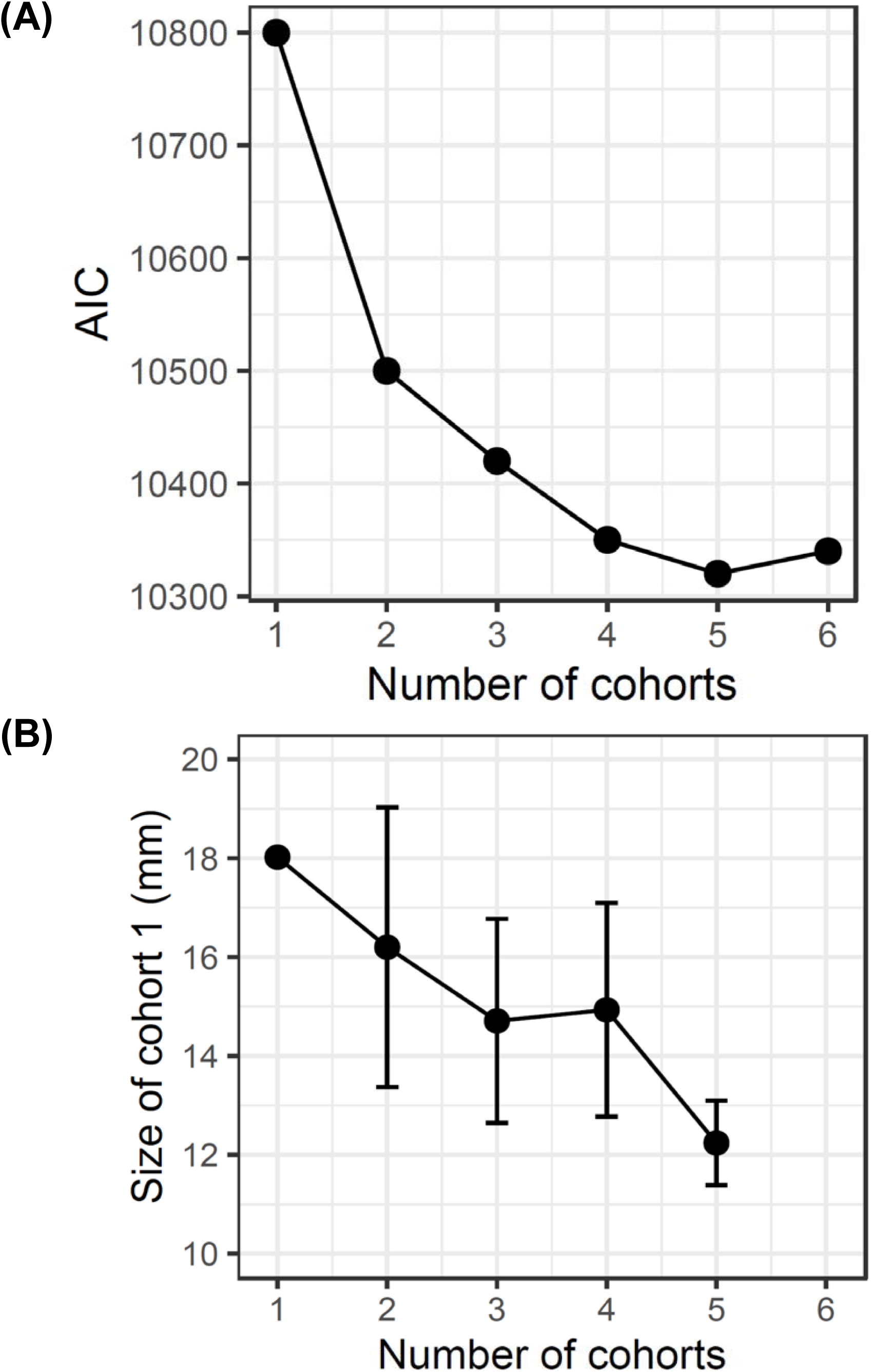
Determination by mixture analysis of the most likely number of zebra mussel cohorts colonizing the experimental panels in Cardiff Bay (undisturbed panels) showing (A) changes in AIC values depending on number of putative cohorts, and (B) mean size (± SD) of the smallest cohort (cohort 1).Table 1. Examples of eradication programmes of zebra mussel.

## References

Adams DC, and Lee DJ. 2012. Technology adoption and mitigation of invasive species damage and risk: application to zebra mussels. Journal of Bioeconomics 14:21–40.

Aldridge DC, Elliott P, and Moggridge GD. 2004. The recent and rapid spread of the zebra mussel (Dreissena polymorpha) in Great Britain. Biological Conservation 119:253–261.

Aldridge DC, Elliott P, and Moggridge GD. 2006. Microencapsulated BioBullets for the control of biofouling zebra mussels. Environmental Science & Technology 40:975–979.

Alix M. 2010. Zebra mussel (*Dreissena polymorpha*) population in the newly formed Cardiff Bay. Cardiff University.

Alix M, Knights R, and Ormerod SJ. 2016. Rapid colonisation of a newly formed lake by zebra mussels and factors affecting juvenile settlement. Management of Biological Invasions 7:405–418.

Allen YC, Thompson BA, and Ramcharan CW. 1999. Growth and mortality rates of the zebra mussel, Dreissena polymorpha, in the Lower Mississippi River. Canadian Journal of Fisheries and Aquatic Sciences 56:748–759.

Baker S, and Hornbach D. 1997. Acute physiological effects of zebra mussel (*Dreissena polymorpha*) infestation on two unionid mussels, Actiononaias ligamentina and Amblema plicata. Canadian Journal of Fisheries and Aquatic Sciences 54:512–519.

Bastviken DT, Caraco NF, and Cole JJ. 1998. Experimental measurements of zebra mussel (*Dreissena polymorpha*) impacts on phytoplankton community composition. Freshwater Biology 39:375–386.

Berkman PA, Haltuch MA, Tichich E, Garton DW, Kennedy GW, Gannon JE, Mackey SD, Fuller JA, and Liebenthal DL. 1998. Zebra mussels invade Lake Erie muds. Nature 393:27.

Bowman MF, and Bailey R. 1998. Upper pH tolerance limit of the zebra mussel (*Dreissena polymorpha*). Canadian Journal of Zoology 76:2119–2123.

Casagrandi R, Mari L, and Gatto M. 2007. Modelling the local dynamics of the zebra mussel (*Dreissena polymorpha*). Freshwater Biology 52:1223–1238.

Chase ME, and Bailey RC. 1999. The ecology of the zebra mussel (*Dreissena polymorpha*) in the lower Great Lakes of North America: I. Population dynamics and growth. Journal of Great Lakes Research 25:107–121.

Churchill CJ, Hoeinghaus DJ, and La Point TW. 2017. Environmental conditions increase growth rates and mortality of zebra mussels (*Dreissena polymorpha*) along the southern invasion front in North America. Biological Invasions 19:2355–2373.

Claudi R, Prescott T, Mastisky S, and Coffey H. 2014. Efficacy of copper based algaecides for control of quagga and zebra mussels. Picton (ON): RNY Consulting Report prepared for California Department of Water Resources, Aquatic Nuisance Species Program.

Cleven E-J, and Frenzel P. 1993. Population dynamics and production of *Dreissena polymorpha* (Pallas) in River Seerhein, the outlet of Lake Constance (Obersee). Archiv für Hydrobiologie 127:395–407.

Costa R, Aldridge D, and Moggridge G. 2011. Preparation and evaluation of biocide-loaded particles to control the biofouling zebra mussel, *Dreissena polymorpha*. Chemical Engineering Research and Design 89:2322–2329.

Decksbach N. 1935. Dreissena polymorpha—Verbreitung im europäischen Teile der UdSSR und die sie bedingenden Faktoren: Mit 1 Karte. Internationale Vereinigung für theoretische und angewandte Limnologie: Verhandlungen 7:432–438.

Durán C, Lanao M, Anadón A, and Touyá V. 2010. Management strategies for the zebra mussel invasion in the Ebro River basin. Aquatic Invasions 5:309–316.

Fahnenstiel GL, Bridgeman TB, Lang GA, McCormick MJ, and Nalepa TF. 1995. Phytoplankton productivity in Saginaw Bay, Lake Huron: effects of zebra mussel (*Dreissena polymorpha*) colonization. Journal of Great Lakes Research 21:464–475.

Fernald RT, and Watson BT. 2013. Eradication of zebra mussels (*Dreissena polymorpha*) from Millbrook quarry, Virginia. Quagga and Zebra Mussels: Biology, Impacts and Control Lewis Publishers, Boca Raton, FL:195–214.

Fong PP, Kyozuka K, Duncan J, Rynkowski S, Mekasha D, and Ram JL. 1995. The effect of salinity and temperature on spawning and fertilization in the zebra mussel *Dreissena polymorpha* (Pallas) from North America. The Biological Bulletin 189:320–329.

Glomski LM. 2015. Zebra Mussel Chemical Control Guide, Version 2.0. Engineer Research and Development Center Vicksburg MS Environmental Lab.

Haag WR, and Garton DW. 1992. Synchronous spawning in a recently established population of the zebra mussel, *Dreissena polymorpha*, in western Lake Erie, USA. Hydrobiologia 234:103–110.

Hammer Ø, Harper DA, and Ryan PD. 2001. PAST: paleontological statistics software package for education and data analysis. Palaeontologia electronica 4:9.

Hincks SS, and Mackie GL. 1997. Effects of pH, calcium, alkalinity, hardness, and chlorophyll on the survival, growth, and reproductive success of zebra mussel (*Dreissena polymorpha*) in Ontario lakes. Canadian Journal of Fisheries and Aquatic Sciences 54:2049–2057.

Holland RE. 1993. Changes in planktonic diatoms and water transparency in Hatchery Bay, Bass Island area, western Lake Erie since the establishment of the zebra mussel. Journal of Great Lakes Research 19:617–624.

Johnson LE, Ricciardi A, and Carlton JT. 2001. Overland dispersal of aquatic invasive species: a risk assessment of transient recreational boating. Ecological Applications 11:1789–1799.

Kassambara A, and Mundt F. 2017. Package ‘factoextra’. CRAN

Kearney M, and Morton B. 1970. The distribution of *Dreissena polymorpha* (Pallas) in Britain. Journal of Conchology 27:o97–100.

Kilgour BW, Mackie GL, Baker MA, and Keppel R. 1994. Effects of salinity on the condition and survival of zebra mussels (*Dreissena polymorpha*). Estuaries 17:385.

Kinzelbach R. 1992. The main features of the phylogeny and dispersal of the zebra mussel. In: Neumann D, Jenner HA (eds) The zebra mussel Dreissena polymorpha: ecology, biological monitoring and first applications in the water quality management. Gustav Fischer, Stuttgart. Limnol Aktuell 4:4–17

Kobak J. 2001. Light, gravity and conspecifics as cues to site selection and attachment behaviour of juvenile and adult *Dreissena polymorpha* Pallas, 1771. Journal of Molluscan Studies 67:183–189.

Kobak J. 2004. Recruitment and small-scale spatial distribution of *Dreissena polymorpha* (Bivalvia) on artificial materials. Archiv für Hydrobiologie 160:25–44.

Lewis DP, and Whitby GE. 1997. Method and apparatus for controlling zebra and related mussels using ultraviolet radiation. Google Patents.

Lovell SJ, Stone SF, and Fernandez L. 2006. The economic impacts of aquatic invasive species: a review of the literature. Agricultural and Resource Economics Review 35:195–208.

Lowe S, Browne M, Boudjelas S, and De Poorter M. 2000. 100 of the world’s worst invasive alien species: a selection from the global invasive species database: Invasive Species Specialist Group Auckland.

Lund K, Cattoor KB, Fieldseth E, Sweet J, and McCartney MA. 2018. Zebra mussel (*Dreissena polymorpha*) eradication efforts in Christmas Lake, Minnesota. Lake and Reservoir Management 34:7–20.

Luoma JA, Severson TJ, Barbour MT, and Wise JK. 2018. Effects of temperature and exposure duration on four potential rapid-response tools for zebra mussel (Dreissena polymorpha) eradication. Management of Biological Invasions 9:425–438.

Luoma JA, Weber KL, Severson TJ, and Mayer DA. 2015. Efficacy of *Pseudomonas fluorescens* strain CL145 A spray dried powder for controlling zebra mussels adhering to native unionid mussels within field enclosures. Reston (VA): US Geological Survey Report 1050.

Mackie GL, and Schloesser DW. 1996. Comparative biology of zebra mussels in Europe and North America: an overview. American Zoologist 36:244–258.

MacNeil C, Platvoet D, Dick JT, Fielding N, Constable A, Hall N, Aldridge D, Renals T, and Diamond M. 2010. The Ponto-Caspian ‘killer shrimp’, *Dikerogammarus villosus* (Sowinsky, 1894), invades the British Isles. Aquatic Invasions 5:441–445.

Mari L, Bertuzzo E, Casagrandi R, Gatto M, Levin SA, Rodriguez□Iturbe I, and Rinaldo A. 2011. Hydrologic controls and anthropogenic drivers of the zebra mussel invasion of the Mississippi□Missouri river system. Water Resources Research 47.

Mari L, Casagrandi R, Bertuzzo E, Rinaldo A, and Gatto M. 2014. Metapopulation persistence and species spread in river networks. Ecology Letters 17:426–434.

Martel A. 1993. Dispersal and recruitment of zebra mussel (*Dreissena polymorpha*) in a nearshore area in west-central Lake Erie: the significance of postmetamorphic drifting. Canadian Journal of Fisheries and Aquatic Sciences 50:3–12.

Martel A, Mathieu AF, Findlay CS, Nepszy SJ, and Leach JH. 1994. Daily settlement rates of the zebra mussel, *Dreissena polymorpha*, on an artificial substrate correlate with veliger abundance. Canadian Journal of Fisheries and Aquatic Sciences 51:856–861.

Minchin D, Lucy F, and Sullivan M. 2002. Zebra mussel: impacts and spread. Invasive aquatic species of Europe Distribution, impacts and management: Springer, 135–146.

Minchin D, and Moriarty C. 1998. Zebra mussels in Ireland.

Nalepa TF, Cavaletto JF, Ford M, Gordon WM, and Wimmer M. 1993. Seasonal and annual variation in weight and biochemical content of the zebra mussel, *Dreissena polymorpha*, in Lake St. Clair. Journal of Great Lakes Research 19:541–552.

Nalepa TF, Wojcik JA, Fanslow DL, and Lang GA. 1995. Initial colonization of the zebra mussel (*Dreissena polymorpha*) in Saginaw Bay, Lake Huron: population recruitment, density, and size structure. Journal of Great Lakes Research 21:417–434.

O’Neill JCR. 1997. Economic impact of zebra mussels-results of the 1995 National Zebra Mussel Information Clearinghouse Study. Great Lakes Research Review 3:35–44.

Pedroli J. 1977. Relation entre les oiseaux aquatiques et Dreissena polymorpha dans le lac de Neuchâtel. Ornithol Beob 74:86–87.

Ram JL, Fong PP, and Garton DW. 1996. Physiological aspects of zebra mussel reproduction: maturation, spawning, and fertilization. American Zoologist 36:326–338.

Ricciardi A, Rasmussen J, and Whoriskey F. 1995. Predicting the intensity and impact of *Dreissena* infestation on native unionid bivalves from Dreissena field density. Canadian Journal of Fisheries and Aquatic Sciences 52:1449–1461.

Rodriguez-Rey M, Consuegra S, Borger L, and Garcia de Leaniz C. under review. Boat ramps facilitate the dispersal of the highly invasive zebra mussel (Dreissena polymorpha). Royal Society Open Science.

Schloesser DW, Nalepa TF, and Mackie GL. 1996. Zebra mussel infestation of unionid bivalves (Unionidae) in North America. American Zoologist 36:300–310.

Seaver RW, Ferguson GW, Gehrmann WH, and Misamore MJ. 2009. Effects of ultraviolet radiation on gametic function during fertilization in zebra mussels (*Dreissena polymorpha*). Journal of Shellfish Research 28:625–634.

Spidle AP, May B, and Mills EL. 1995. Limits to tolerance of temperature and salinity in the quagga mussel (*Dreissena bugensis*) and the zebra mussel (*Dreissena polymorpha*). Canadian Journal of Fisheries and Aquatic Sciences 52:2108–2119.

Stanczykowska A, and Lewandowski K. 1993. Thirty years of studies of *Dreissena polymorpha* ecology in Mazurian Lakes of Northeastern Poland. IN: Zebra Mussels: Biology, Impacts, and Control Lewis Publishers, Boca Raton, FL 1993 p 3-37, 17 fig, 8 tab, 83 ref, 4 append.

Strayer DL. 2010. Alien species in fresh waters: ecological effects, interactions with other stressors, and prospects for the future. Freshwater Biology 55:152–174.

Strayer DL, D’Antonio CM, Essl F, Fowler MS, Geist J, Hilt S, Jarić I, Jöhnk K, Jones CG, and Lambin X. 2017. BoomLbust dynamics in biological invasions: towards an improved application of the concept. Ecology Letters 20:1337–1350.

Vanhaecke D, Garcia de Leaniz C, Gajardo G, Thomas C, and Consuegra S. 2012. Metapopulation dynamics of a diadromous galaxiid fish and potential effects of salmonid aquaculture. Freshwater Biology 57:1241–1252.

Waller DL, and Bartsch MR. 2018. Use of carbon dioxide in zebra mussel (*Dreissena polymorpha*) control and safety to a native freshwater mussel (Fatmucket, *Lampsilis siliquoidea*). Management of Biological Invasions 9:439–450.

Walz N. 1978. The energy balance of the freshwater mussel *Dreissena polymorpha Pallas in laboratory experiments and in Lake Constance. II*. Reproduction Arch Hydrobiol.

Watters A, Gerstenberger SL, and Wong WH. 2013. Effectiveness of EarthTec® for killing invasive quagga mussels (*Dreissena rostriformis bugensis*) and preventing their colonization in the Western United States. Biofouling 29:21–28.

Whitledge GW, Weber MM, DeMartini J, Oldenburg J, Roberts D, Link C, Rackl SM, Rude NP, Yung AJ, and Bock LR. 2015. An evaluation Zequanox® efficacy and application strategies for targeted control of zebra mussels in shallow-water habitats in lakes. Management of Biological Invasions 6:71–82.

Wimbush J, Frischer ME, Zarzynski JW, and Nierzwicki□Bauer SA. 2009. Eradication of colonizing populations of zebra mussels (*Dreissena polymorpha*) by early detection and SCUBA removal: Lake George, NY. Aquatic Conservation: Marine and Freshwater Ecosystems 19:703–713.

Wisniewski R. 1974. Distribution and character of shoals of *Dreissena polymorpha* Pall. In the Bay part of Goplo Lake. Prace Limnology 8:73–81.

Wood C, Bishop J, and Yunnie A. 2015. Comprehensive reassessment of NNS in Welsh marinas.

Wood SN. 2001. mgcv: GAMs and generalized ridge regression for R. R news 1:20–25.

